# Quantifying the Influence of the Actin Cytoskeleton on Ion Transport in Dendritic Spines by Homogenization of the Poisson-Nernst-Planck Equations

**DOI:** 10.1101/2023.08.03.551796

**Authors:** Florian Eberhardt

## Abstract

Dendritic spines are filled with a very dense actin cytoskeleton. However, due to their small size, the impact of this mesh on biophysical parameters has not been studied so far, and it remains unclear to what extent it might affect ion flow in dendritic spines. Only recently has the three-dimensional internal structure of dendritic spines been quantified in great detail in electron microscopic tomography data. Based on these results, we estimate the effect of the spine actin cytoskeleton on diffusion and permittivity. We apply a method called homogenization to estimate effective diffusion tensors and permittivity tensors in Poisson-Nernst-Planck (PNP) equations. We find that the volume taken up by the intracellular structure alone cannot explain the changes in these biophysical parameters. The characteristic architecture of the intracellular space in dendritic spines will reduce the diffusion of ions by 33% to 46% and the permittivity by 30% to 42%, compared to values found for the cytosol free of intracellular structures.

These results can be used to improve computational studies using PNP equations and help to better interpret experimental results of electrical and chemical compartmentalization.

## 1 Introduction

Dendritic spines harbor almost all excitatory synapses on pyramidal cells [29]. They typically have a distinctive mushroom shape, characterized by a head and a neck [5], and are filled with a dense actin cytoskeleton [10]. However, the cytoskeleton in dendritic spines differs from the structures found in other cell domains, such as lamellipodia, which contain parallel and long bundles of actin [38]. In dendritic spines, the cytoskeleton is a highly branched and extremely dense mesh, with branching points occurring every few nanometers. The filaments and associated proteins form a uniform, dense mesh that fills the entire spine [15].

The increased density of the intracellular structure affects the diffusion of molecules and ions in dendritic spines. For example, it has been shown that the presence of actin-related proteins in spine necks can alter the chemical compartmentalization of spines [40]. However, the structure and composition of the intracellular space not only affect the diffusion of molecules [30], but also the migration of ions and, therefore, the electrical resistance in spines [11].

To study dendritic spines, computer simulations based on Poisson-Nernst-Planck (PNP) equations are often used [12, 13, 19, 24]. These equations involve biophysical parameters such as diffusion and permittivity, which must be accurately known to model dendritic spines. However, experimental measurement of these parameters is challenging due to the small size of spines [41]. Consequently, the parameters used in simulations of spines are often based on measurements from other cell domains or even other cell types. It remains an open question how exactly the intracellular structure affects ionic currents in spines.

In this work, we theoretically estimate the effect of a specific organization of the intracellular space on the parameters of diffusion and permittivity in dendritic spines. The studied geometries are based on the results of a quantitative analysis of the intracellular space in dendritic spines from electron microscopic data [15, 23]. To estimate the effective values of diffusion and permittivity, we use a method called homogenization. The homogenization procedure replaces a microscopically heterogeneous medium (the intracellular space) with a homogeneous approximation [4, 6]. In the case of PNP equations, this will result in effective diffusion and permittivity parameters [17, 36]. This method has already been applied to study ionic transfer in cementitious materials [7], viscous fluids in bituminous concretes [9], and diffusion of molecules in biological cells [30].

In dendritic spines, the actin cytoskeleton and associated proteins reduce the volume available for the diffusion and migration of ions. We found that for parallel bundles of filaments, the reduction of diffusion and permittivity values along the orientation of the filaments can be explained by the excluded volume effect. However, for geometries that form a dense mesh, such as the actin cytoskeleton in spines, the excluded volume effect alone cannot explain the reduction in diffusion and permittivity. In [15], it was found that the available volume is reduced by 20% to 30% in hippocampal pyramidal cells (average of 28%). Based on these numbers, we computed effective tensors and found that this would reduce the diffusion of ions in the intracellular space by 33% to 46%, and the permittivity would be reduced by 30% to 42%. Therefore, the particular organization of the intracellular space is important in estimating biophysical parameters in dendritic spines.

In the future these results will help to improve the accuracy of computer simulations of dendritic spines and help in better interpreting experimental results.

## 2 Methods

### Homogenization of the PNP-equations

Homogenization is a method used to approximate a heterogeneous microscopic structure with a homogeneous one. This enables the estimation of macroscopic values for diffusion and permittivity in the PNP-equations from a microscopically heterogeneous intracellular space. The PNP-equations have been previously utilized to study electrodiffusion in dendritic spines [12, 24].

In the following, we will summarize the homogenized PNP-equations based on the formalism presented by Schmuck and Bazant [36]. The intracellular space of spines is filled with a dense actin cytoskeleton and other molecules (shown as dark regions in Fig. 1A,B), as well as an electrolyte, the cytosol (shown as light regions in Fig. 1A,B). It is assumed that the intracellular space can be subdivided into two separate domains, a solid phase (cytoskeleton and macromolecules) and a liquid phase (electrolyte). We assume that the electrolyte contains ions such as *Na*^+^, *K* ^+^, and *Cl*^−^ with negligible size. Diffusion and migration of ions only occur in the electrolyte, but the electric field penetrates both domains. There are no surface reactions, and no-flux boundaries at the solid-liquid interface are applied for ion flow. For the electric potential, it is continuous at the solid-liquid interface. Negative surface charges on cytoskeletal filaments and larger molecules with low diffusivities are modeled as a continuous fixed background charge. The diffusion constant has a fixed value in the liquid phase and is zero in the solid phase. The permittivity has two distinct positive values in the liquid and solid phase on the microscale.

**Figure 1:**
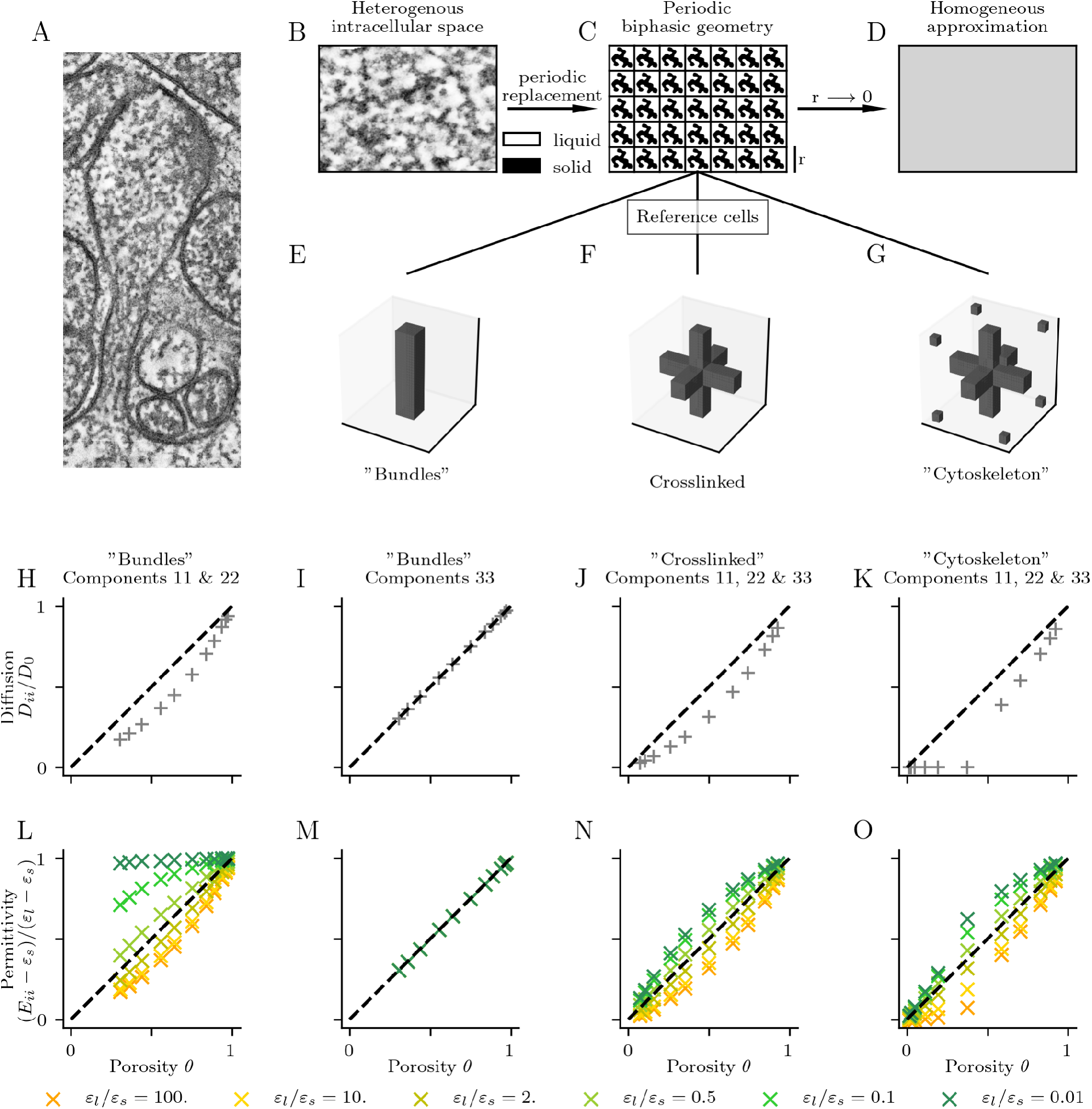
Homogenization of the intracellular space in spines. A) Electron tomography data of a hippocampal pyramidal cell spine that was analyzed in [15]. B) Heterogeneous intracellular space with stained filaments and proteins (dark regions) and fluid cytosol (light regions). C) The real structure gets replaced by a periodic arrangement of reference cells. D) Homogenized PNP-equations with effective diffusion and permittivity tensors are obtained by solving the reference cell problem. The reference cell problem can be derived by letting the size of the reference cell approach zero. Different reference cells represent parallel bundles of filaments (E), cross-linked filaments (F), and a dense filamentous mesh found in spines (G). Panels H-O show the diagonal elements of the diffusion and permittivity correction tensors. Porosity represents the relative volume taken by the liquid phase. The permittivity correction tensors were computed for different ratios between the permittivities of the liquid and solid phase.

It was found that the organization of the cytoskeleton is uniform throughout the intracellular space of dendritic spines, with nodes in the mesh occurring every 15 nm [15]. Hence, the length scale of the mesh is small compared to the size of spines, which are several hundred nanometers. This confirms the assumption that microscopic material properties vary rapidly, at a length scale *l*, compared to the macroscopic size of the sample *L*. Specifically, this means that *r* = *l/L <<* 1.

To apply homogenization, the intracellular structure is replaced by a periodic tiling of reference cells with a characteristic length *l* (Fig. 1C). A reference cell must contain the relevant properties of the intracellular organization, and, in addition to [36], we assume local thermodynamic equilibrium at the length scale *l*. As the size of the reference cell is decreased, while the sample size *L* and the volume fraction (solid/liquid) are held constant, a new set of equations is obtained in the limit *r*→ 0. Solving these equations finally results in the effective diffusion and permittivity tensors of the PNP-equations.

For a comprehensive introduction to homogenization, we recommend [14]. An intuitive explanation and experimental validation were presented by Auriault et al. [6]. An overview of different homogenization techniques, such as two-scale convergence and asymptotic expansion, can be found in [4, 34]. Both techniques have been applied to derive effective equations for PoissonNernst-Planck systems [8, 17, 34, 35] or more complicated Stokes-Poisson-Nernst-Planck systems [31, 33, 36]. Numerical applications can be found, for example, in [7, 8, 16, 30].

### Dimensionless PNP-equations

In this paragraph we summuarize the relevant results derived by Schmuck and Bazant [36]. First we define dimensionless variables. All dimensionless variables are denoted by a “∼”. Moreover we assume a binary electrolyte with charge numbers *z*_1_ = +1 and *z*_2_ = −1.

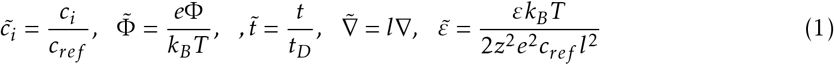

*c*_*i*_ denotes the concentration of ion-species *i* and *c*_*ref*_ denotes a reference salt concentration. Φ is the electric potential, *e* the elementary charge, *k*_*B*_ the Boltzmann constant and *T* the temperature. Moreover *l* is the reference length scale and *t*_*D*_ = *l*^2^*/D* the reference time scale. Finally *D* is the diffusion constant on the microscale. Altogether this leads to the following dimensionless PNPequations:

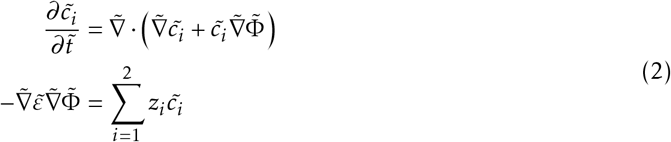

According to equation (1.8) from [36] we can write the permittivity as

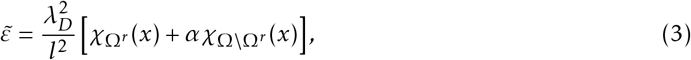

where *ε*_*l*_ and *ε*_*s*_ are the permittivity values of the liquid and solid domain on the microscale and 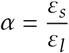 is a constant. The Debye length in the electrolyte is 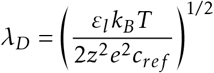 . Below, always the dimensionless variables are used, but to simplify the notation the “∼” on the variables is dropped again.

Effective diffusion and permittivity tensors Schmuck and Bazant [36] use an asymptotic multiscale expansion to derive macrocopic PNP-equations, where *x* represents the macroscale, *y* = *x/r* the microscale and *r* = *l/L* denotes the scale parameter. The scalar variables 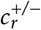 and Φ *r*are expanded using the ansatz 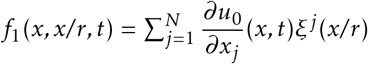 for the first order term. The periodic functions *ξ*^*j*^ are corrector functions with a period equal to the size of the reference cell *Y* = [0, *l*_1_] × [0, *l*_2_] × … × [0, *l*_*n*_]. In particular, the expansion to the first order reads as

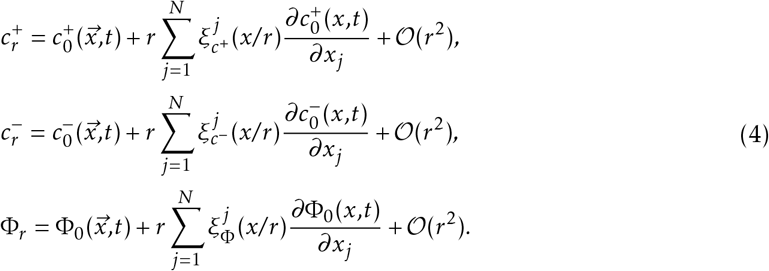

Inserting eq. (4) into eq. (2) and sorting the resulting equations by powers of *r* leads to three sets of equations of order 𝒪 (*r*^−2^), 𝒪 (*r*^−1^), and 𝒪 (1). (Remark: We will not consider the migration-tensor, and Stokes-flow will be neglected, i.e., the fluid velocity is set to zero, compared to [36]).

Based on [36], from equations of order 𝒪 (*r*^−2^) one can deduce that 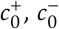 and Φ_0_ are constant on the microscale (constant on a reference cell). That implies that 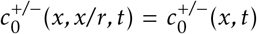 and 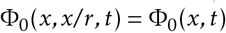 as already indicated in eq. (4).

From equations of order 𝒪 (*r*^−1^), one can derive the reference cell problems, that uniquely define the corrector functions 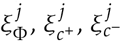 The notation of the reference cell problems was taken from [33], but the Einstein sum convention was taken out and replaced by gradient and divergence operators as in equation (5) of [34]. Then the reference cell problem corresponding to the electric potential reads as

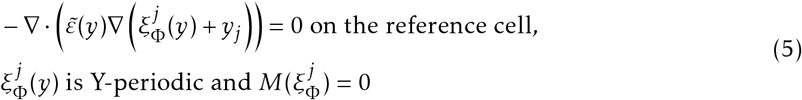

and for the concentration variables the reference cell problem is

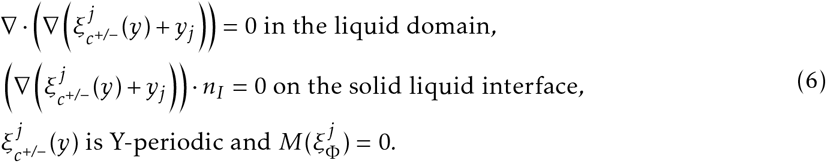

Here, 𝕄 (.) denotes the mean value on the reference cell *Y*, and *j* = 1, …, *N* denotes the spatial dimension of the reference cell (e.g., *N* = 3). *n*_*I*_ is the normal vector on the solid-liquid interface. An explanation of the function spaces of the corrector functions *ξ* can be found in equation (1.12) of [36].

Next, by collecting equations of (1), one obtains the effective diffusion-dispersion correction tensor 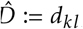 and the effective electric permittivity tensor 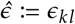, With 1 ≪*K,l* ≪ *N*.

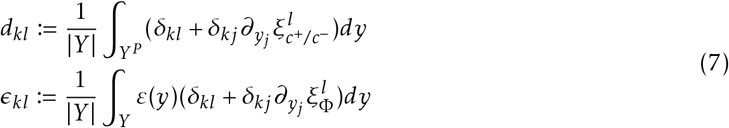

Here *δ* denotes the Kronecker-*δ* and *∂*_*yl*_ denotes the partial derivative with respect to *y*_*j*_ . *Y*^*p*^ ⊂ *Y* is the volume taken by the solid domain (Fig. 1C) of the reference cell. Furthermore, the porousity Θ is defined as

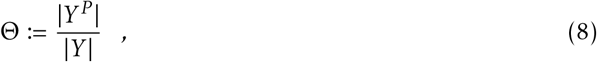

and the surface charge density per unit volume *ρ*_*s*_ is computed by

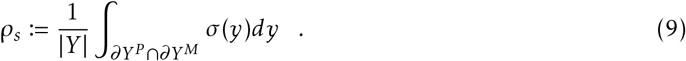

The surface charge on the solid-liquid interface becomes a continuous background charge. Finally, the dimensionless upscaled PNP-system can be stated:

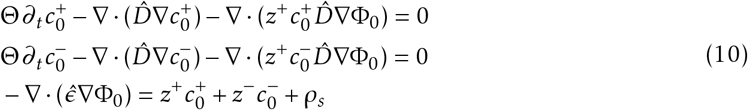

Numerical implementation The effective diffusion-dispersion and electric permittivity tensors are computed using the FEniCS [2, 27] computing platform with DOLFIN v. 2018.1.0 [1, 25]. FEniCS is a sophisticated open-source platform for solving partial differential equations with the finite element method. This computing platform uses several components such as FFC [3, 22, 42], UFL [28], and FIAT [20, 21].

To implement in DOLFIN, the problem consisting of differential equations and boundary conditions has to be cast into its weak form. For practical reasons, we implement the boundary condition𝕄 (*ξ*) = 0 not in the function space of test and trial functions, but with a Lagrange multiplier *λ*. In the weak form the corrector functions *ξ* have to fullfill the following equation,

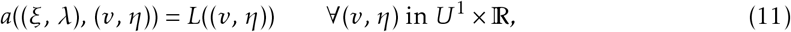

where *a*((*u, λ*), (*v, η*)) is called the bilinear form and *L*((*v, η*)) is called the linear form. The solution lies in the function space

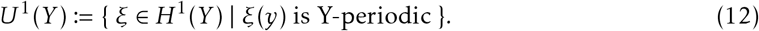

*H*^1^ is the standard notation for the Sobolev space.

Now the bilinear form and linear form of the weak problem can be defined for the corrector functions *ξ* as

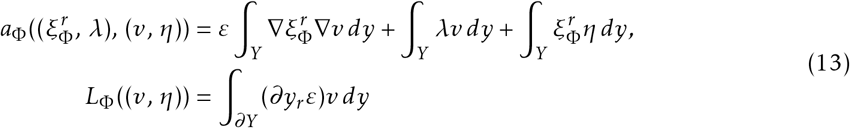

and

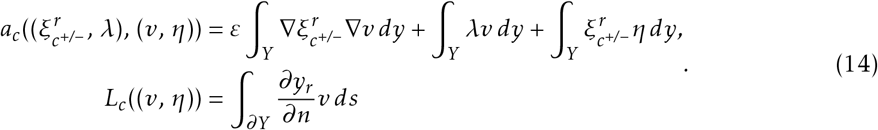

Here, *y*_*r*_ is r-*th* component of 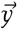*ds* denotes the integral over the boundary of the liquid domain.

Next, we introduce a triangulation 𝒯_*h*_ of Y into triangles/tetrahedrons (2D/3D) *K*_*i*_ and the finite dimensional subspace

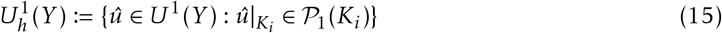

Effectively, we create an equispaced mesh of triangles (2D) or tetrahedrons (3D) with P_1_ Lagrange finite elements [26]. Finally, we find solutions in FeniCS for 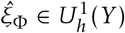 and 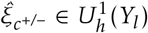 that fullfill the following weak forms of equations 6 and 5

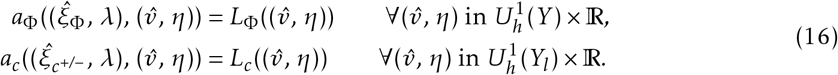

## 3 Results

In this study, we used a method called homogenization to replace the heterogeneous intracellular space of dendritic spines with a homogeneous approximation with effective diffusion and permittivity values. The homogenization was applied to the Poisson-Nernst-Planck equations, as presented by Schmuck and Bazant [36].

To estimate the effective diffusion-dispersion and permittivity tensors, we replaced the heterogeneous mesh of filaments (Fig. 1A,B) with a periodic arrangement of reference cells (Fig. 1C). The reference cell must be characteristic of the geometry of the structure, with the same length scales or porosity (relative volume taken by the liquid phase). The reference cell also consists of a solid phase (molecules) and a liquid phase (electrolyte). Diffusion and drift of ions, such as *Na*^+^, *K* ^+^, and *Cl*^−^, occur only in the liquid phase, whereas the electric field penetrates both subdomains. To homogenize the system, we decreased the size of the reference cell *l*, while holding the sample size *L* and the volume fraction (solid/liquid) constant. In the limit *r* →0, with *r* = *l/L*, we obtained a set of equations whose solution finally led to the effective diffusion and permittivity tensors (Fig. 1B-C).

### Convergence of the solution

To verify our implementation, we first reproduce the diffusion tensors found in [6]. A sketch of the experimental setup and the reference cell can be seen in Fig. 2A. Auriault and Lewandowska [6] found a value of *D*11 = 0:3833 for the normalized diffusion correction tensor (normalized with respect to the porosity). The equations were solved with 144 finite elements in [6]. This is in agreement with our results at low resolution (376 triangles). Experimentally, [6] found 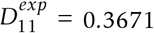 which is comparable to our result of 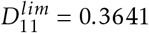 in the case of a very fine mesh.

**Figure 2:**
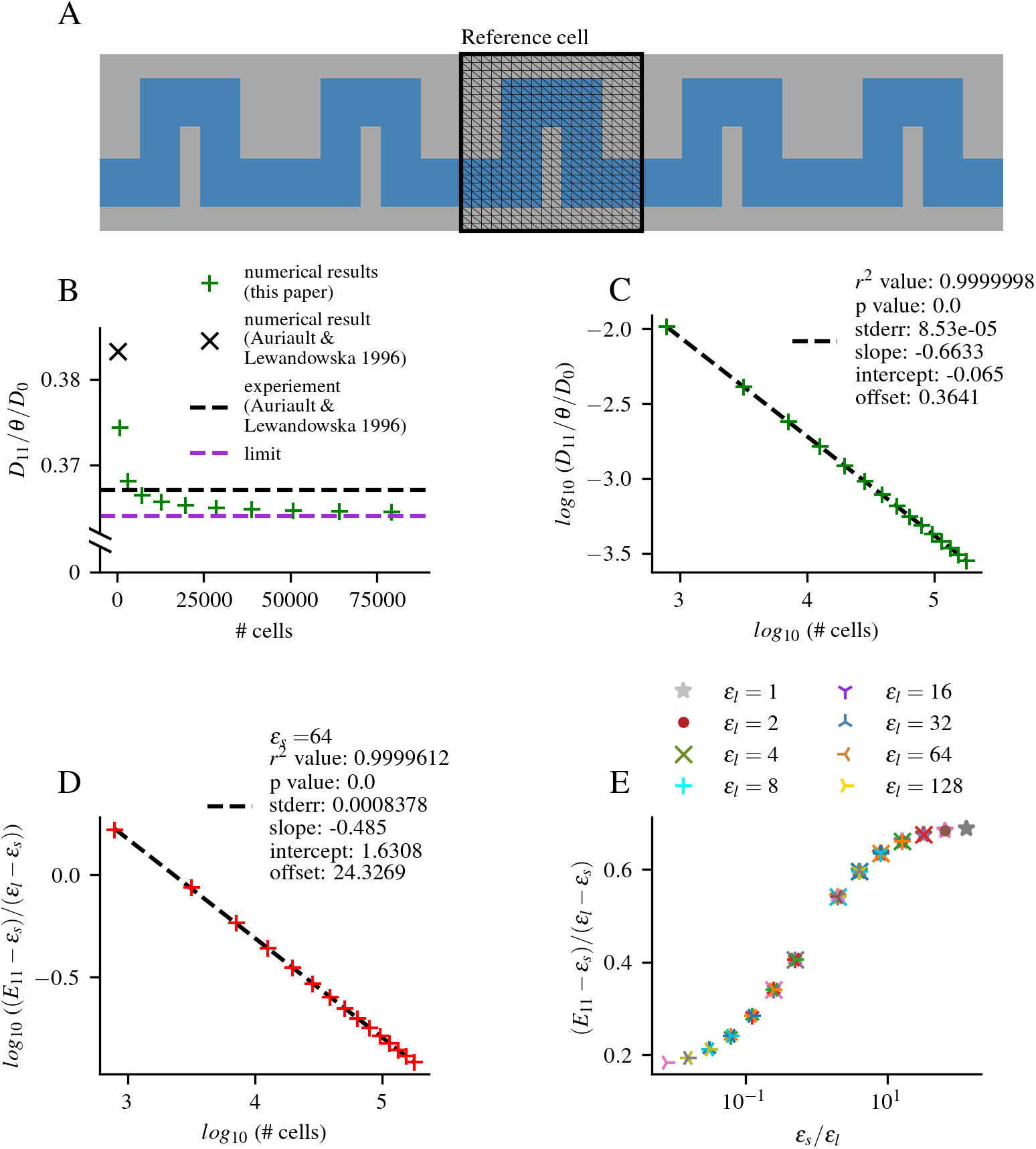
Verification and convergence of the results. A) The setup and reference cell studied in [6]. B) The numerically estimated values of the normalized diffusion correction tensor converge close to the experimental result. C) After subtracting the limit value (purple dotted line in B), the convergence can be explained by a power law, as seen in linear behavior in a log-log plot. By optimizing the explained variance, the limit (purple dotted line) can be estimated. D) This procedure is applied to all components of the diffusion and permittivity tensors. E) The normalized permittivity correction tensor only depends on the ratio between the permittivities in the solid and liquid domains.

In general, the accuracy of a solution improves for finer meshes and converges to the exact solution for an infinite number of elements. However, the computational complexity, memory consumption, and runtime increase drastically for refined meshes [26]. This is especially important for 3D geometries, where the number of elements and the computational load increase with the third power of the mesh resolution. Therefore, we study the convergence of the correction tensors as a function of the number of triangles or cells in the mesh. We use triangular meshes for all 2D geometries and tetrahedral meshes for all 3D geometries.

As expected, the values of the elements in the correction tensors are always converging to a limit value as a function of the number of finite elements (Fig. 2B). The convergence of all entries in the correction tensors *C*_*ij*_ as a function of the number of triangles/tetrahedrons *n* can always be very accurately described by a power function of the form 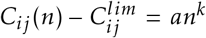 (where *a* and *k* are some constants). The limit value 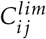 is found by optimizing the explained variance of a linear regression fit in a log-log plot (Fig. 2C,D). This is also true for geometries that cannot be exactly represented by triangular elements on a regular mesh (e.g., round geometries), and the explained variance was 99.7% on average for all studied 2D and 3D reference cells. This allowed us to accurately estimate the correction tensor even for 3D geometries where the resolution was limited. All tensor elements shown in Figs. 1H-O, 2E, 4, and 5C,D are therefore the estimated limit values.

Finally, the normalized permittivity tensor elements (*E*_*ij*_ −*ε*_*s*_)*/*(*ε*_*l*_− *ε*_*s*_) do not depend on the absolute values of *ε*_*s*_ and *ε*_*l*_, but only on the ratio between the two permittivities of the solid and liquid domain. To verify this, we alter both permittivities between 1, 2, 4, 8, 16, 32, 64, 128 on the *Perturbed channel 2D* reference cell (Fig. 3A), and for each combination, we compute the correction tensors (Fig. 2E).”

**Figure 3:**
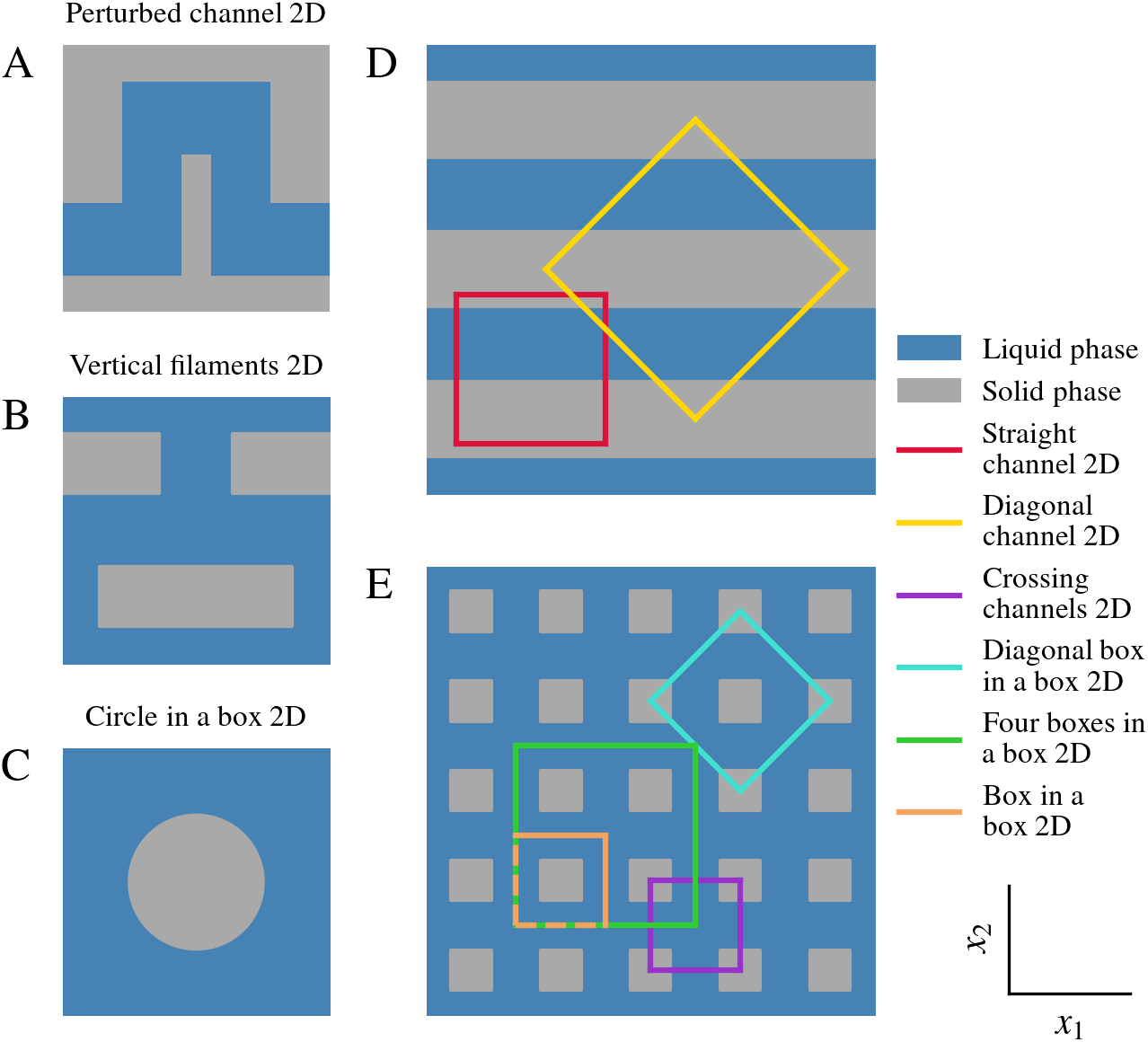
2D reference cells. We compute correction tensors for various 2D reference cells. For some geometries, such as D) and E), different reference cells are possible. Each reference cell has its own name, as shown on the right.

### Homogenization of various 2D-Geometries

To further test the numerical implementation and to study the effect of different structures on diffusion and permittivity, we compute the correction tensors for various 2D geometries (Fig. 3A-E).

In the simple case of the *Straight channel 2D* reference cell (Fig. 3D), the correction tensors can even be computed analytically. The tensor component for diffusion in the *x*_1_-direction is determined by the porosity of the material, *D*_11_ = Θ. As there is no migration of ions possible in the *x*_2_-direction, it follows that *D*_22_ = 0 (compare [36]). To analytically compute the permittivity correction tensors for the *Straight channel 2D* reference cell, we consider an ideal plate capacitor filled with alternating layers of two different electrolytes. This is identical to a periodic arrangement of the reference cells as shown in Fig. 3D. Homogenization of this structure now means passing the size of the reference cell to zero while keeping the volume constant.

To begin with, we assume *n* periodic reference cells in the *x*_1_ direction and *n* periodic reference cells in the *x*_2_ direction. In this case, the liquid domain has permittivity *ε*_*l*_ = 2 and the solid domain has *ε*_*s*_ = 1. Both domains, liquid and solid, have the same volume.

To compute *E*_11_, the volume can be considered as *n* parallel capacitors, while to compute *E*_22_, the volume can be considered as *n* capacitors in a row. The off-diagonal elements are zero. Here, *l*_*x*_ = *l*_*y*_ = *l* denotes the size of the reference cells and *L*_*x*_ = *L*_*y*_ = *L* denotes the size of the full domain. Moreover, we extend the domain into the *x*_3_ direction by a length of *l* and *L*, respectively. The capacitor surface is given by *A* = *L*^2^, and the distance between the capacitor plates is given by *d* = *L*. Finally, we define the scale parameter as the number of reference cells *n* = *L/l*. We find the values in the permittivity correction tensors by taking the limit *n* → ∞:

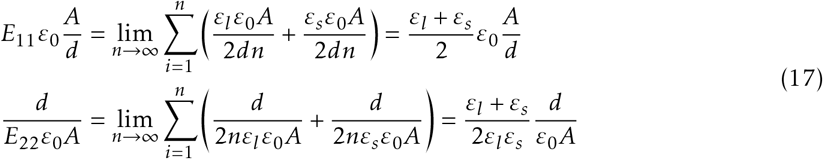

The numerical results in Figure 4 exactly match the theoretically predicted values *E*_11_ = 1.5 and *E*_22_ = 1.333 for a porosity Θ = 0.5 and *ε*_*l*_*/ε*_*s*_ = 2.

**Figure 4:**
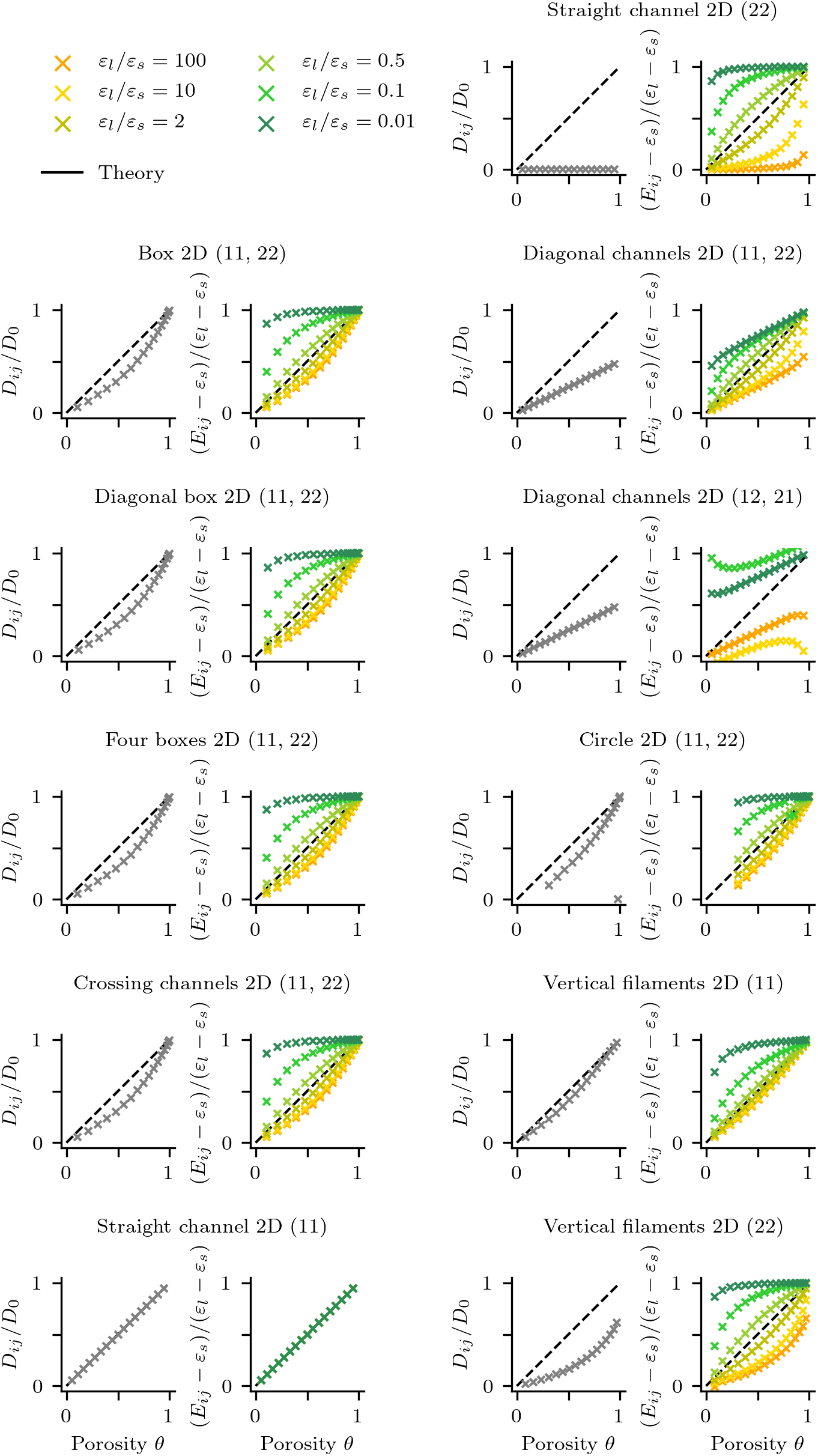
Correction tensors for 2D geometries. The effective diffusion and permittivity correction tensors for the 2D reference cells shown in Figure 3.

For some structures, different reference cells can be used to represent the microscopic geometry. However, the correction tensors should not depend on the choice of the particular reference cell. This assumption is confirmed by the finding that the three reference cells, *Box 2D, Crossing channels 2D*, and *Four boxes 2D* (Fig. 3), all lead to the same correction tensors (Fig. 4).

Next we study the effect of rotations. A rotation of the *Box 2D* reference cell by *π/*4 leads to the *Diagonal box 2D* reference cell. The numerical results can be seen in Fig. 3. The same rotation can also be applied to the correction tensors instead of the reference cell. Therefore, we transform the correction tensors of the *Box 2D* reference cell by the corresponding rotation matrix *ε*^∗^ = *R*(*α*)*εR*^−1^(*α*). In fact, both approaches lead to identical results. As the non-diagonal elements are zero and the two diagonal elements are identical, the structure is isotropic and the tensors can be reduced to constants.

Furthermore, we rotate the *Straight channel 2D* reference cell to obtain the *Diagonal channels 2D* reference cell. As before, we compute the correction tensors in two ways: numerically from the rotated reference cell and by applying the rotation matrix to the correction tensors. Once again, the results of both approaches are identical. However, in this case, the correction tensors of the rotated reference cell contain non-vanishing diagonal elements, indicating that the structure is not isotropic.

To reliably implement the periodic boundary conditions on the reference cell, we use finite elements whose nodes lie on a regular triangular grid. At lower grid resolutions, especially in 3D, it is therefore only possible to accurately implement geometries that consist of rectangles or cuboids (Fig. 1E-G). But intuitively, one would rather represent an actin filament by a cylinder. Therefore, we compare the difference in the effective diffusion tensors for round and square solid domains in 2D (Fig. 5A,B). At a high resolution, a good approximation of the geometries can be achieved. We find that for the same porosity, round and square geometries lead to similar correction tensors (Fig. 5C). The relative difference between the diagonal elements of tensors is always lower than 4.2 % (Fig. 5D).”

**Figure 5:**
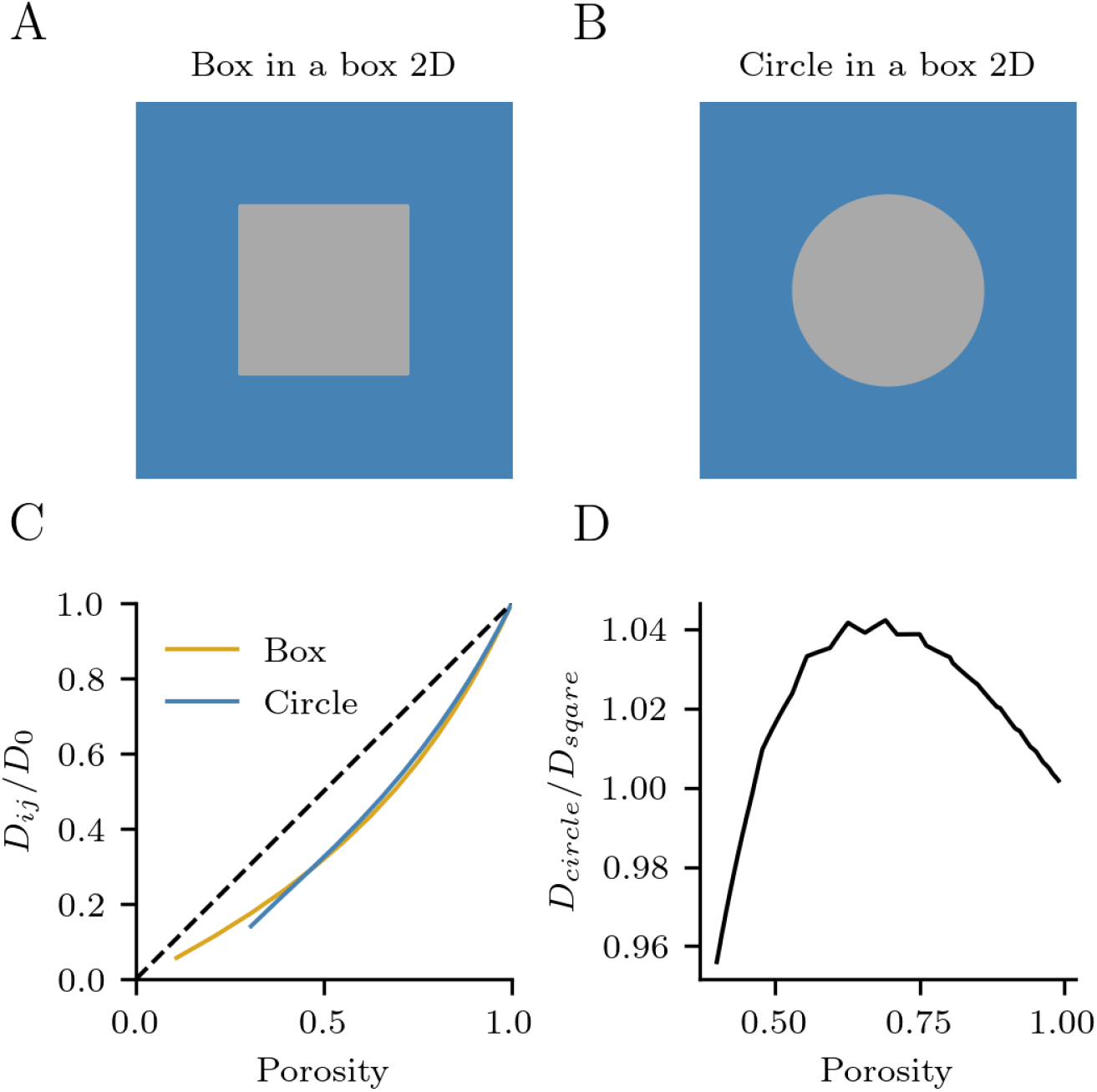
Error of square geometries. The filaments in the 3D geometries are represented by elongated rectangular shapes. The maximum relative difference between the round and square geometries is 4.2 %. Therefore, approximating round filaments by rectangles only produces small errors.

### Diffusion and Permittivity are Decreased in Dendritic Spines

In the final section, we estimate diffusion and permittivity correction tensors for three different reference cells in 3D. The first reference cell (Fig. 1E) represents an intracellular space that is filled by parallel bundles of F-actin or microtubules. The second reference cell (Fig. 1F) represents a structure of strongly crosslinked filaments. The third reference cell (Fig. 1G) contains a combination of strongly crosslinked filaments and fragments of filaments or other proteins and represents the intracellular organization in dendritic spines. All solid domains carry a negative surface charge.

We find that, along the elongated structures in the *Bundles* reference cell (Fig. 1E), the diffusion tensor component *D*_33_ is only reduced by the porosity. However, perpendicular to the orientation of the bundles, the tensor components are reduced, compared to the porosity, and *D*_11_ and *D*_22_ are identical to the values found for the 2D reference cell *Box in a box 2D* (Fig. 2E, Fig. 3).

For the *Crosslinked* and *Cytoskeleton* reference cells, the diffusion and normalized permittivity correction tensors (in the case *ε*_*l*_*/ε*_*s*_≥ 10) are well below the porosity. Therefore, porosity alone cannot explain the values of the components in correction tensors. For porosities Θ *>* 0.5, the reduction in diffusion and permittivity is similar for both geometries. However, for very low porosities, the effective diffusion suddenly drops in the case of the *Cytoskeleton* reference cell. This is consistent with the fact that for low porosities, the liquid domain consists of disconnected subdomains, making diffusion impossible.

The cytoskeleton geometry (Fig. 1F,G) is isotropic, as all three diagonal elements have the same value and all off-diagonal elements are vanishing. In this case, the diffusion tensor can be replaced by a constant again. Based on previous results, the porosity in hippocampal pyramidal cell spines is between 0.7 and 0.8 [15], and for the permittivity, it can be assumed that 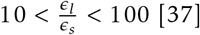

This allows us to estimate the correction tensors in dendritic spines to be 0.56 *< D*^*ef f*^ */D*_0_ *<* 0.67 and 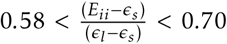. From this, we can conclude that the actin cytoskeleton significantly reduces diffusion and permittivity in dendritic spines.

## 4. Discussion

The size of dendritic spines is too small to reliably measure many biophysical parameters in experiments [41]. However, an accurate knowledge of these parameters is needed for the interpretation of experimental results [39] and for computational models of dendritic spines [24]. In this paper, we estimate the diffusion constant and permittivity in dendritic spines based on the intracellular organization in a mathematically rigorous way. Therefore, we apply a method called homogenization to the Poisson-Nernst-Planck equations, which is a set of partial differential equations. The theory was previously presented by Schmuck and Bazant [36]. Homogenization of the PNP equations allows us to compute effective diffusion and permittivity tensors for a microscopically heterogeneous material. It replaces the composite material by a homogeneous approximation (Fig. 1A-D). Each individual microscopic structure leads to a characteristic set of biophysical constants [16] on the macro-scale. We estimate the diffusion and permittivity correction tensors for various different geometries in 2D (Fig. 3) and 3D (Fig. 1E-G). In this way, we accurately verify the method (Fig. 2) and provide an intuitive overview of how a specific structure affects the diffusion and permittivity (Fig. 4). Finally, we study the effect of the cytoskeleton in dendritic spines (Fig. 1K,O).

The intracellular space of dendritic spines is filled with a dense meshwork of filaments and proteins. Based on our findings in [15], we assume that the porosity (the volume occupied by the liquid electrolyte domain) in dendritic spines lies between 0.7 and 0.8. In a reference cell that mimics the intracellular space of spines (Fig. 1G), we find that the diffusion constant of ions will reduce between 33% and 46% (Fig. 1K) compared to its value in the cytosol (free of intracellular structures). The permittivity will reduce accordingly from 33% to 42% (Fig. 1O) compared to a cytosol without intracellular structures.

In dilute solutions, the diffusion of sodium, potassium, and chloride was measured as 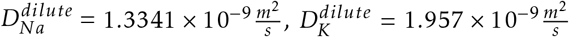 and 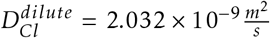 respectively [32]. The macromolecular content of the cytosol reduces the diffusion coefficient in the cytosol without intracellular structures by approximately 20% compared to the diffusion in dilute solutions [30]. The cytoskeleton and related proteins further reduce the diffusion by a multiplicative factor. These values can be found in the diffusion correction tensors *D*_*ij*_ studied in this work. Finally, one can estimate the diffusion constants of ions in dendritic spines (compare with [30]) as 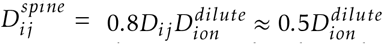. These numbers are valid in the spine head and along the main axis of the spine neck, where the overall size is significantly larger than that of a reference cell [5, 33].

In this work, we studied the effect of a characteristic intracellular spine geometry on biophysical parameters. However, it is also likely that differences between individual spines on a neuron exist. For example, it was found that the diffusional coupling between spine head and dendrite changes by a factor of up to 10 in spines undergoing long-term potentiation [18]. This would imply a shrinkage of the spine neck diameter by a factor of three. Therefore, it is likely that the structure of the intracellular space was affected as well. Using super-resolution light microscopy, another study found that the presence of synaptopodin alters the structure of the neck cytoskeleton [40], and thereby alters the diffusion of mGluR5 receptors in the spine neck. We think that in the future, the combination of electron microscopy, light microscopy, and homogenization can help to connect macroscopic parameters of individual spines to biochemical processes related to the spine cytoskeleton.

## Acknowledgements

I am deeply grateful to my professor and supervisor, Prof. Dr. Andreas V. M. Herz, for his guidance and support throughout my doctoral studies. Furthermore, I would like to acknowledge the generous support of the Ludwig-Maximilians Universität, the German Federal Ministry of Education and Research, and the Bernstein Center for Computational Neuroscience Munich, who provided the funding necessary to carry out this research.

